# Identification of Successful Mentoring Communities using Network-based Analysis of Mentor-Mentee Relationships across Nobel Laureates

**DOI:** 10.1101/075432

**Authors:** Julia H. Chariker, Yihang Zhang, John R. Pani, Eric C. Rouchka

**Affiliations:** Department of Psychological and Brain Sciences, Life Sciences Building, Room 317, University of Louisville, Louisville, KY, 40292, USA.; KBRIN Bioinformatics Core, 522 East Gray Street, University of Louisville, Louisville, KY, 40202, USA.; Department of Computer Engineering and Computer Science, Duthie Center for Engineering, Room 208, University of Louisville, Louisville, KY, 40292, USA.

## Abstract

Skills underlying scientific innovation and discovery generally develop within an academic community, often beginning with a graduate mentor’s laboratory. In this paper, a network analysis of doctoral student-dissertation advisor relationships in The Academic Tree is used to identify successful mentoring communities in high-level science, as measured by number of Nobel laureates within the community. Nobel laureates form a distinct group in the network with greater numbers of Nobel laureate ancestors, descendants, mentees/grandmentees, and local academic family. Subnetworks composed entirely of Nobel laureates extend across as many as four generations. Successful historical mentoring communities were identified centering around Cambridge University in the latter 19^th^ century and Columbia University in the early 20^th^ century. The current practice of building web-based academic networks, extended to include a wider variety of measures of academic success, would allow for the identification of modern successful scientific communities and should be promoted.

## Background

High achievement in intellectual innovation has been measured in part with the awarding of prestigious honors, such as the Nobel Prize. From the first awards in 1901 through the awards in 2015, a total of 573 prizes have been awarded to 900 laureates, including 875 individuals and 25 organizations ^1^. To some extent, the scientific knowledge and skill underlying these achievements has been transmitted across generations through person-to-person academic mentoring, and much attention has been given to individual mentoring relationships^2–6^. However, interaction within a scientific laboratory extends beyond the mentor-mentee relationship. Laboratories make up a community of researchers in which knowledge and skill is shared within and across generations through relationships between academic siblings and between mentees and their grandmentors. Within the realm of high-level science, successful mentoring communities can provide clues to creating fertile environments for scientific innovation.

The contribution of mentoring to academic success is difficult to isolate within an entire scientific population because additional factors, such as level of institutional resources and student talent, also vary across training situations. Successful researchers generally attract more federal and institutional resources and more talented students, and success begets success. However, when looking for successful mentors and mentoring communities within a subset of high achievers, such as Nobel laureates, these factors should operate somewhat equally. With modern technology and the benefits of crowd-sourcing, generations of mentoring relationships are now represented in networks such as The Academic Tree ^7,8^, greatly facilitating the study of mentoring on a large scale. The Academic Tree is a vast crowd-sourced network containing mentor-mentee relationships across several interconnected domains of science. If Nobel laureates are a distinctive group due in some way to mentoring communities, greater connectedness should be found among them in the network. Otherwise, we should find Nobel laureates randomly dispersed across the network. If Nobel laureates are a distinctive group, and quality of mentoring plays an important role in their success, it should be possible to identify particularly successful mentoring communities within the group of laureates (i.e., the “best of the best”).

We looked for connectedness among Nobel laureates in The Academic Tree by asking whether they have a greater number of Nobel laureate academic family members than non-Nobel laureates have. We restricted our analysis to doctoral student-advisor relationships and assessed academic family structure in several ways. We examined the number of Nobel laureate ancestors for each individual as well as the number of local and global descendants. Local descendents covered two generations in the network and included mentees and grandmentees, whereas global descendants comprised all generations of descendants. To identify more dispersed mentoring communities, we looked at the number of Nobel laureates within each individual’s local academic family, including three generations in all directions in the network. Three generations encompassed an individual’s mentor, grandmentor, great-grandmentor, mentees, grandmentees, great-grandmentees, sibling, aunts, and uncles. We compared the outcomes of this analysis to results obtained from many topologically identical networks in which Nobel status was randomly assigned across all individuals in each of the networks.

Nobel laureates appear to be a distinct group with a greater number of Nobel laureate family members than non-Nobel laureates have on all measures. In addition, several historical scientific communities exist with high concentrations of Nobel laureates. In some instances, Nobel laureates are directly connected to one another over three and four generations of scientists. Biographical and historical accounts offer the only access to characteristics associated with these successful communities. However, with the expansion of current network databases to include a variety of performance measures for all scientists, new methods could be used to identify modern scientific communities and to study them more directly.

## Results

### Descriptive Summary.

As can be seen in Table 1, the distributions for the number of academic family members and the number of Nobel laureate academic family members are positively skewed on all measures. On some measures, the range was quite large, prompting a closer look at The Academic Tree. For example, the range for number of descendants extended to 2,628 for Nobel laureates and 13,620 for non-Nobel laureates. However, academic lineages have been recorded across several centuries, justifying these numbers. For example, Michele Savonarola, a non-Nobel laureate physician scientist, practicing in the 15^th^ century, has 13,620 descendants, 73 Nobel descendants, and 0 ancestors in the network. Wilhelm Friedrich Ostwald, a Nobel Prize winner in chemistry (1909), has five ancestors and the highest number of descendants in the Nobel laureate group at 2,628. In terms of mentees/grandmentees, it appears that Robert B. Woodward, a Nobel Prize winner in chemistry (1965), has 213 mentees/grandmentees recorded, one of whom is a Nobel laureate. In the non-Nobel laureate group, Gilbert Stork, a Professor of Chemistry Emeritus at Columbia University, has 149 mentees/grandmentees, also with one Nobel laureate among them. The range for number of local academic family was also quite large. Robert Woodward, a Nobel laureate in chemistry (1965), has the largest local family with 558 members, 5 of whom are Nobel laureates. Of the non-Nobel laureates, Robert T. Paine, professor emeritus of zoology at The University of Washington has 446 local family. The distributions for all measures are displayed in Fig. S1.

**Table 1.** The range and median for number of Nobel laureate academic family and total number of academic family across all measures for Nobel laureates (NL) and non-Nobel laureates (Non-NL). The correlation between number of academic family and number of Nobel laureate academic family for each measure is also displayed. Ancestors refer to individuals moving backward in the directed network, and descendants are all individuals moving forward in the network. M/GM refers to the number of mentees and grandmentees (two generations forward), and local family refers to the number of individuals within 3 generations forward and backward in the network.

|  | Number of<br>NL Academic Family |  | Number of<br>Academic Family |  | Corr. Between<br>Academic Family<br>and NL Academic<br>Family |
| --- | --- | --- | --- | --- | --- |
|  | NL | Non-NL | NL | Non-NL |  |
|  | Range(Mdn) | Range(Mdn) | Range(Mdn) | Range(Mdn) | Spearman's *r* |
| Ancestors | 0 - 6 (0) | 0 - 8 (0) | 0 - 75 (7.5) | 0 - 131 (9) | 0.33*** |
| Descendants | 0 - 21 (0) | 0 - 73 (0) | 0 - 2,628 (11) | 0 - 13,620 (0) | 0.24*** |
| M/GM | 0 - 8 (0) | 0 - 8 (0) | 0 - 213(5) | 0 - 149 (0) | 0.17*** |
| Local Family | 0 - 18 (2) | 0 - 17 (0) | 3 - 558 (31) | 3 - 446 (22) | 0.17*** |

\*\*\* $p < 0.0001$

One reason a strong positive skew was found for ancestors, descendants, and mentees/grandmentees was due to the nature of network data, where some individuals serve as source nodes without ancestors and other individuals serve as sink nodes without descendants.This increases the number of outcomes measuring zero in the data. In this case, having zero Nobel family members is a result of having zero family members. Alternatively, a number of individuals in the network have ancestors or descendants, but none of them are Nobel laureates. Clearly, there are two possible sources for zero Nobel family members. In the analysis of ancestors who were Nobel laureates, for example, there were 2,890 individuals with no ancestors, and thus no Nobel laureate ancestors. On the other hand, there were 37,608 individuals with ancestors, none of whom were Nobel laureates. Similarly, in the analysis of descendants who were Nobel laureates, there were 40,044 individuals having no descendants. At the same time, there were 16,869 individuals with immediate descendants and 17,197 individuals with mentees/grandmentees, none of whom were Nobel Prize winners.

### Approach to Analysis.

Zero-inflated regression models were developed for analyzing data with two possible sources for zero outcomes. With this approach, two models are estimated, a zero-inflation model and a count model ^9,10^. The zero-inflation model is estimated first, using a binomial model to estimate the probability of excess zeros in the data (i.e., a zero outcome due to the absence of family). Once this probability is estimated, the probability for the remaining outcomes is estimated using a Poisson or negative binomial model, whichever is appropriate. In the current paper, zero-inflated models were used to control for excess zeros in estimating the number of Nobel laureate ancestors, descendants, and mentees/grandmentees. There was no need for this in estimating the number of local Nobel laureate family, because inclusion in a connected network necessarily meant that at least one family connection existed.

For all four analyses, negative binomial models were chosen to adjust for greater than expected dispersion in the data (i.e., a high variance to mean ratio). Spearman’s correlations (see Table 1) indicated that the number of Nobel laureate family members was positively related to the size of the academic family. Therefore, in each case, the size of the academic family was entered along with Nobel status as a predictor of the size of the Nobel laureate academic family. As described in the method, the significance level for each analysis was adjusted by comparing the observed test statistics with a distribution of expected test statistics, derived from 1,000 topologically identical networks, each with a random permutation of Nobel status. The regression model coefficients and the distributions of random coefficients used to adjust the significance levels of predictors in the models are available in Table S1 and Fig. S2, respectively.

### Regression Model Outcomes.

Nobel laureates had a greater number of Nobel laureate ancestors than non-Nobel laureates did, suggesting that Nobel laureate mentorship may play a role in the development of future Nobel Prize winners (adjusted *p* = 0.003). However, the number of academic ancestors was not a significant predictor of the number of Nobel ancestors (adjusted *p* = 0.389). Similarly, Nobel laureates had a greater number of Nobel laureate descendants than non-Nobel laureates did (adjusted *p* < 0.001) with number of descendants not significantly predicting number of Nobel laureate descendants (*p* = 0.143).

In contrast to the previous two results, the number of mentees/grandmentees did serve as a significant predictor of number of Nobel laureate mentees/grandmentees (adjusted *p* < 0.001). Still, after controlling for family size, Nobel laureates had a greater number of Nobel laureate mentees and grandmentees than did non-Nobel laureates (adjusted *p* < 0.001). Finally, Nobel laureates also had a greater number of local Nobel Laureates in their academic family than did non-Nobel laureates (adjusted *p* < 0.001). The number of local academic family members did not significantly predict the number of Nobel laureates (adjusted *p* < 0.964).

### Identification of Nobel Laureate Communities.

To identify highly successful scientific communities, the largest component of The Academic Tree Network, displayed in Fig. 1A, was filtered to include only individuals at or above the 99^th^ percentile for the number of local Nobel laureate family members (99^th^ percentile = 4) and the number of Nobel laureate descendants (99^th^ percentile = 1), along with their first neighbors in the network. This produced one large subnetwork of 1276 individuals and 5 smaller subnetworks ranging in size from 3 to 68 individuals (Fig. 1B). This network remained quite large, and in Fig. 2, first neighbors were removed, producing a more tractable set of 30 subnetworks for analysis, ranging from 1 to 73 individuals. Nobel laureates in the surrounding academic family who contributed to the scores of these individuals are not pictured. Consequently, these subnetworks only display individuals at the center of the local academic family, making the scale of the two largest subnetworks remarkable. A list of individuals in this group, along with the number of family and Nobel laureate family on all measures, is available in Dataset S1. To explore the connectivity among these scientists, high resolution images of Figs. 1B and 2 are available in the supplement with scientist’s names (see Figs. S3 and S4).

**Fig. 1.**
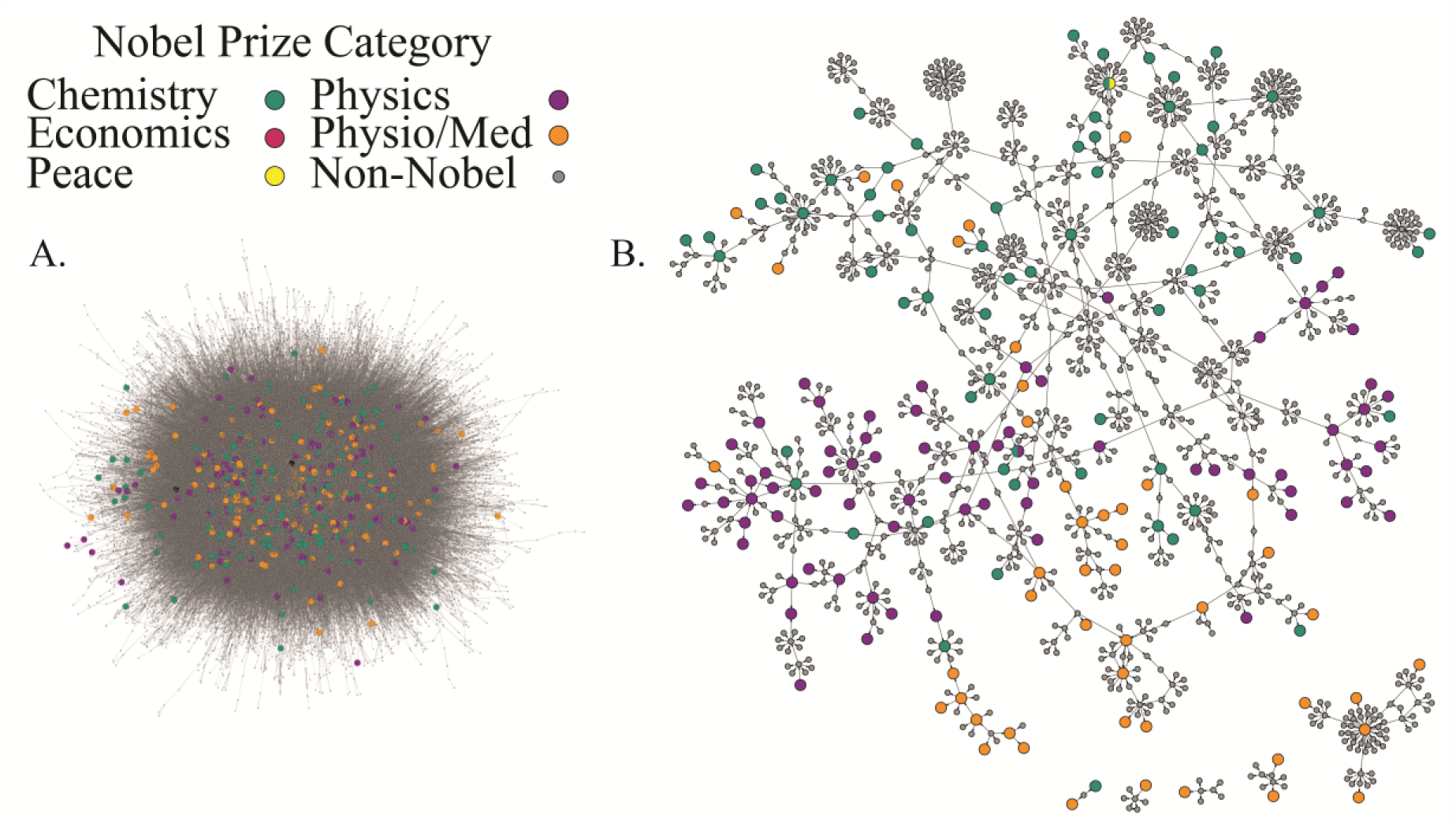
The largest component of The Academic Tree network (A) filtered to include individuals at the 99^th^ percentile for number of Nobel laureate descendants and number of local Nobel family along with their first neighbors (B). The individual names associated with each node in subnetwork B are viewable in a high resolution pdf in the supplement (Fig. S3).

**Fig 2.**
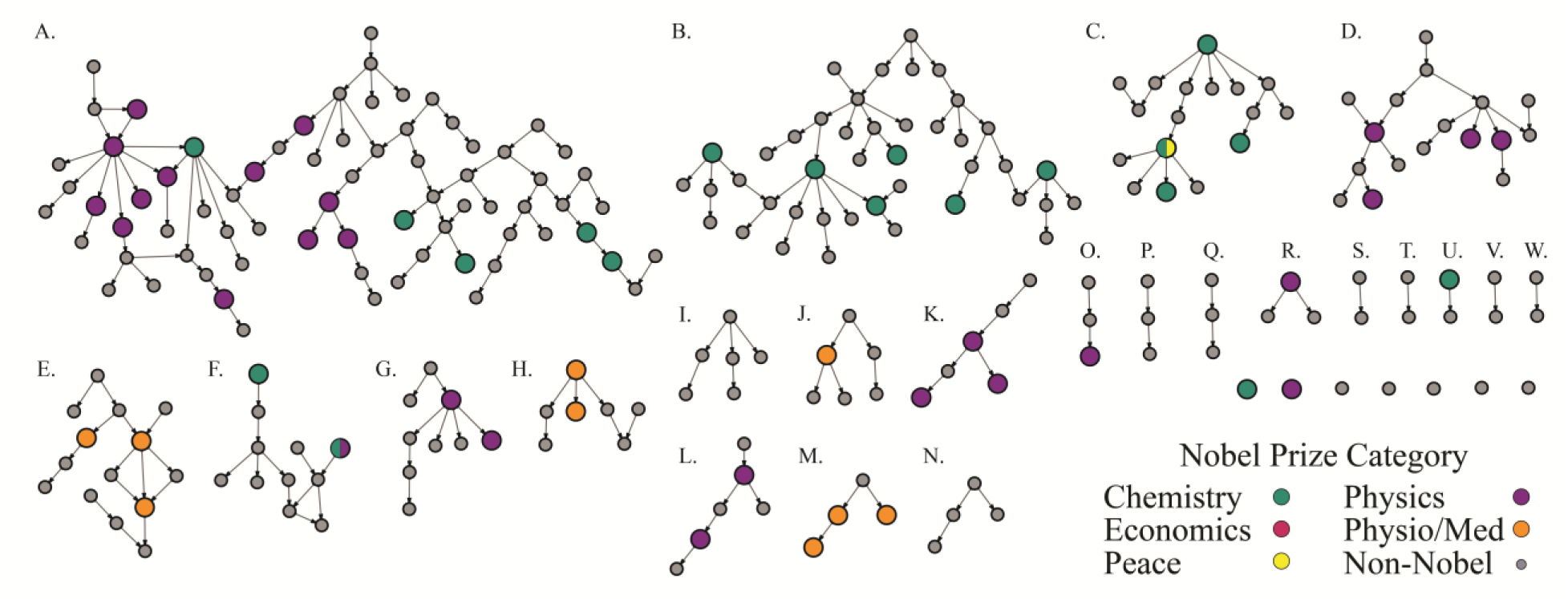
The largest component of The Academic Tree network filtered to include only individuals at the 99^th^ percentile for number of Nobel laureate descendants and number of local Nobel family. The individual names, the number of local Nobel family, and the number of Nobel descendants associated with each node are viewable in a high resolution pdf in the supplement (Fig. S4).

The largest subnetwork (Fig. 2A) can be segmented into two early communities by identifying the geographical location of the scientists. One community centered around J. J. Thomson (physics, 1906) and Ernest Rutherford (chemistry, 1908) at Cambridge University, and a second centered around notable scientists such as August Kundt, Wilhem Rontgen (physics, 1901), Johannes Muller, and Hermann von Helmholtz, among others, working across multiple universities in Germany and Switzerland. The German/Swiss community extends to Herman Staudinger (chemistry, 1953) and Leopold Ruzicka (chemistry, 1939) at the right of the subnetwork. Justis Von Liebig, a German chemist considered the founder of organic chemistry ^11^, has the greatest number of Nobel descendants in this group at 53. Eilhard Mitscherlich, Heinrich Magnus, and Johannes Muller follow with 27, 26, and 26 Nobel descendants, respectively. In the Cambridge community, William Hopkins and Edward Routh, well-known non-Nobel laureate mentors ^12^, lead with 22 Nobel descendants. Their mentees/grandmentees, J.J. Thomson and Ernest Rutherford, both Nobel laureates, have 16 local Nobel laureate family members. David Shoenberg, a British physicist with 17 Nobel laureate family members connects these two major communities.

Interestingly, two additional communities in the largest subnetwork were established in the United States through mentors trained in Germany. Nobel laureate Isador Isaac Rabi (physics, 1944) with 8 Nobel descendants, 6 of which are mentees/grandmentees, serves as the center of one group at Columbia University. William Giauque (chemistry, 1949) and Willard Libby (chemistry, 1960) are at the center of a second community at the University of California, Berkeley.

A significant portion of the second largest subnetwork (Fig. 2B), also contains individuals operating across universities in Germany, and once again, individuals trained in Germany began new communities at universities in Britain, through William Perkins, and universities in the Northeastern United States, through Ira Remsen. In this subnetwork, Friedrich Wohler, a German chemist, has the highest number of Nobel descendants at 39. Johannes Wislicenus, a German chemist, and William Perkin, an English chemist, have the highest number of local Nobel family at 13. Approximately half of Wislicenus’s Nobel family are mentees/grandmentees.

Of particular note in the smaller subnetworks is Enrico Fermi (physics, 1938; Fig. 2K) with 18 Nobel laureate family members and 6 Nobel laureate mentees/grandmentees. Fermi trained and spent his early years as an academic in Italy during the early 20^th^ century, but traveled to Germany to study with Max Born (physics, 1954) and Paul Ehrenfest ^11^. Eventually, near the beginning of WWII, and on winning his Nobel Prize, Fermi moved to the United States and joined the Columbia University community centered on Isaac Rabi. In another subnetwork operating around the same time period (Fig. 2D), Max Born, Werner Heisenberg (physics, 1932), Hans Bethe (physics, 1967), and Robert Oppenheimer, among others, can be found. In this network, Arnold Sommerfeld, a non-Nobel laureate mentor to Heisenberg and Bethe, has 16 Nobel family members and 11 Nobel descendants.

### Nobel Laureate Subnetworks.

On close inspection of Fig. 1B, small subnetworks can be identified that are comprised entirely of Nobel laureates. To explore this further, all non-Nobel laureates were removed from the large strongly connected network (Fig. 1A). Of the 402 Nobel laureates, 260 had no direct connection to another Nobel laureate. However, there were 142 Nobel laureates in 55 subnetworks ranging in size from 2 to 10 individuals. Fig. 3 displays the six largest of the subnetworks. An investigation of relationships in these subnetworks identified seven additional connections recorded in Academic Tree after receiving the data used in the analysis. These are indicated by bold edges in the figure. Once again, the scientific communities at Cambridge and Columbia are identified as exceptional with 13 Nobel laureates connected over four generations at Cambridge (Fig. 3F) and 10 Nobel laureates connected over three generation at Columbia (Fig. 3E).

**Fig. 3.**
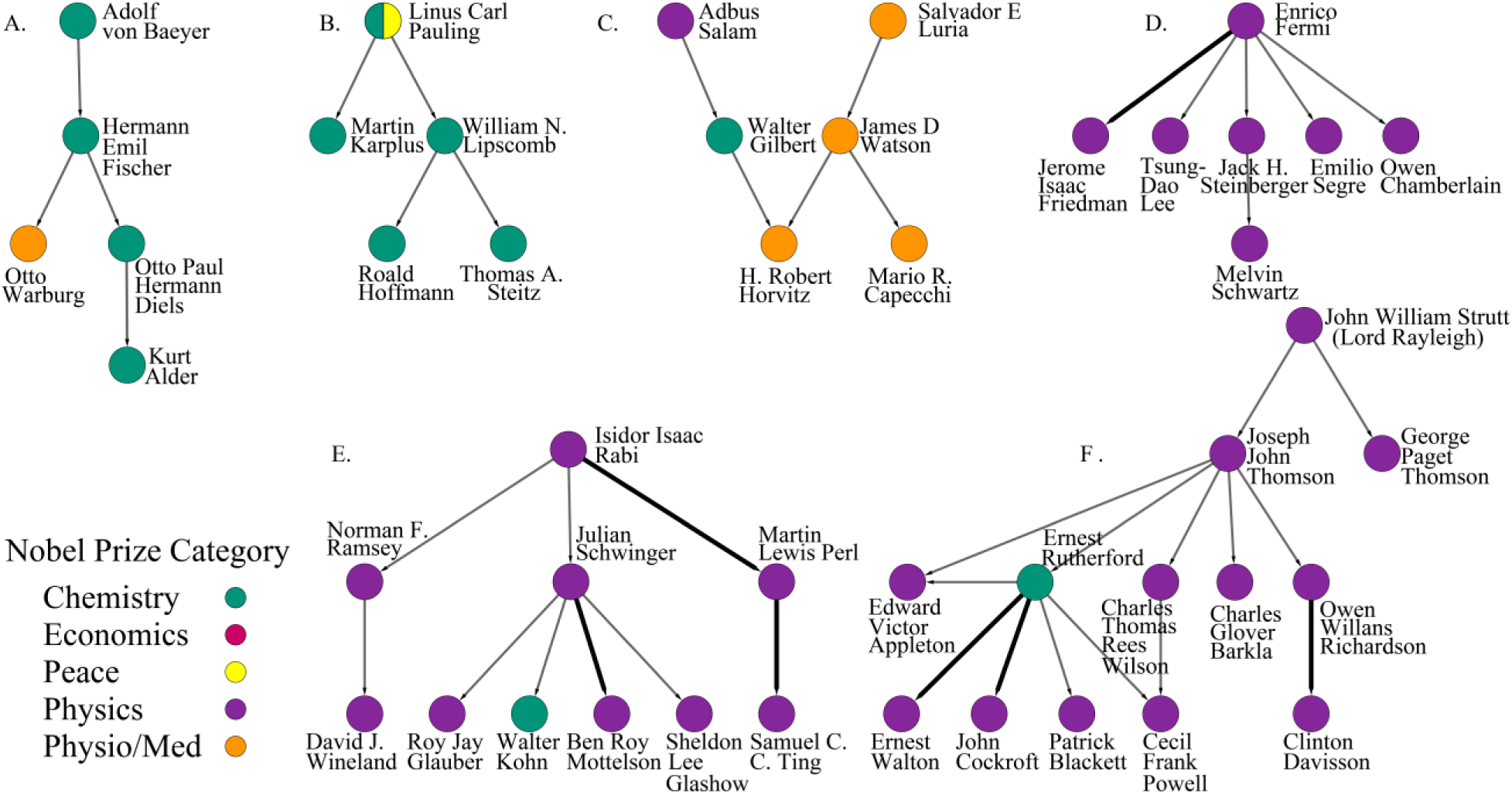
The six largest subnetworks composed entirely of Nobel laureates. Bold edges indicate mentoring relationships recorded in The Academic Tree subsequent to the receipt of the dataset.

### Heterogeneity of Local Academic Family.

In Fig. 1B, clusters of Nobel laureates who were awarded prizes in the same category can be seen with areas of greater diversity appearing where the clusters overlap. In an effort to characterize diversity in the network around Nobel Prize winners, the heterogeneity of Nobel Prize categories was measured within each individual’s local family (see Method).

The vast majority of individuals in the network had 1 or fewer Nobel laureates in their local family. Therefore, the analysis was restricted to individuals with 2 or more Nobel laureates in the family and where some opportunity for diversity existed (205 of 402 Nobel laureates; 3,647 of 57,429 non-Nobel laureates). As the number of Nobel laureates increased, heterogeneity scores also increased (*r* = 0.25, *p* < .0001). The distribution of scores for Nobel Prize winners in individual categories was positively skewed (chemistry: 0 to 0.37, Mdn = 0; physics 0 to 0.42, Mdn = 0.17; Physio/Med: 0 to 0.44, Mdn = 0) and reflective of generally homogeneous clusters of Nobel laureates with some diversity where clusters overlap. There were 6 Nobel laureates (3 physics, 2 physio/med, 1 physics/chemistry) and 39 non-Nobel laureates with scores at or above the 99^th^ percentile (0.37). Archibald Hill, a Nobel laureate in physiology and medicine (1922; Fig. 2E), scored the highest family heterogeneity (0.44) with 4 Nobel laureate family in physiology/medicine, 4 in physics, and 1 in chemistry.

These findings prompted us to ask whether there was any difference in family diversity for Nobel laureates and non-Nobel laureates, and the data were fit to a quasibinomial model with Nobel status and number of local Nobel laureate family as predictors. In the final analysis, heterogeneity scores were not predicted by Nobel status (adjusted *p* = 0.347) after controlling for the number of Nobel laureates in the local family (adjusted *p* = 0.177).

## Discussion

Remarkable connectedness among Nobel laureates is found through generations of mentoring relationships in The Academic Tree network. Nobel laureates have more Nobel laureate ancestors, more local and global descendants, and more local academic family members than do non-Nobel laureates. A variety of explanations for this connectedness exist. Nobel laureates undoubtedly possess superior knowledge and skill that individuals in the local academic family, and the greater community, may acquire through a variety of means. Other factors related to the availability of resources and the attraction of talent are no doubt significant contributors to the connectedness of this group. These additional factors are difficult to separate from the transfer of knowledge through mentoring but are an integral part of any successful scientific community and should be valued as such.

Several areas of the network, representing mentoring relationships in historical scientific communities, were identified with high concentrations of Nobel laureates. In some locations, direct connections between Nobel laureates occurred over three and four generations. When exploring biographical and historical accounts of these communities, it was apparent that much greater interconnectedness existed among scientific communities than is reflected by doctoral mentor-mentee relationships. A high degree of interaction occurred throughout these communities. For, example, after completing a dissertation at Columbia University, Isador Isaac Rabi (physics, 1944) spent over a year in Europe where he encountered some of the greatest minds in science, many of whom went on to become Nobel laureates ^13^. Rabi returned to Columbia to eventually lead the Physics Department and to become a central figure in one of the two most successful communities identified in this analysis ^14–16^. In future work, extending the analysis to include a greater variety of mentoring relationships would better capture the true interconnectivity among scientists.

It is significant that many of the successful communities identified by this network analysis existed at a time when travel and communication were much more difficult than they are today. Ernest Rutherford (chemistry, 1908) traveled from New Zealand to attend Cambridge as one of the first students admitted from outside the university ^17,18^. This occurred in the latter half of the 19th century prior to the invention of the airplane and intercontinental telephone service.At this point in history, physical proximity was critical to the transmission of ideas and expertise. In modern science, however, virtual meetings, video lectures, online courses, and online databases (e.g., PubMed ^19^, Google Scholar ^20^) provide remarkably easy access to current, innovative ideas in science. It seems likely that the mentoring patterns among scientists are being radically altered by greater accessibility to information and each other. Still, for many scientists, it is difficult to imagine that virtual proximity could ever be a satisfying replacement for the day-to-day personal interaction found in a positive mentoring relationship.

Biographical and historical accounts provided the sole access to more detailed information about the communities identified in this study. Warwicke (2003) offers a particularly valuable and compelling account of the scientific community identified at Cambridge in the latter part of the 19^th^ century ^12^. However, modern scientific communities could be studied if the types of data required to identify them were available. Although the number of Nobel laureates within an academic community serves as a legitimate measure of success, especially when the research focus is restricted to high-level science, much more could be accomplished if a variety of other performance measures were readily available and reliably accurate Information regarding publications, impact factors, citations, funding sources, and other awards, would allow for a more sensitive evaluation of success within a community. This could be achieved by a committed effort in the scientific community to collect performance measures from all individuals and universities and to make them available in an open-source database, something The Academic Tree is currently attempting to accomplish.

Several factors would be critical to the success of this endeavor. Primarily, a comprehensive list of all researchers’ publications would need to be available in a centralized, open-source database. Currently, no one source is guaranteed to have a complete set of publications for an individual author ^21^, and publication information must be obtained from multiple sources, such as Web of Science ^22^, Scopus ^23^, and PubMed ^19^. Furthermore, some of these sources are proprietary and require a fee for use. Google Scholar ^20^ has access to several proprietary sources through licensing agreements but does not allow automated searches of its website, something that is a requirement when conducting an analysis of “big data”. As an example, the largest component of the Academic Tree Network analyzed in this study contained 57,831 individuals, making manual search costly in terms of time.

Making a wide variety of performance measures accessible would also increase the value of a database for evaluating scientific success. Number of publications, a measure of productivity, is not a sufficient measure of success. Rather, number of citations, considered a measure of quality, is often factored alongside number of publications in calculations such as the *h*-index ^24^. Number of citations is not consistently available in the sources mentioned earlier, and it is not clear how often this information is updated. Along these lines, additional quality measures, such as a journal’s impact factor at the time of an article’s publication, author funding, and additional awards, would be useful in developing new algorithms for measuring the quality of research and the impact of an individual’s and a community’s contribution to science.

Another critical element in developing an effective database involves the assignment of unique identifiers for scientists. This is especially important when dealing with crowd-sourced data. On one hand, crowd-sourcing allows for the collection of data that would be difficult or impossible to obtain otherwise. On the other hand, a quick glance at the Academic Tree dataset makes it clear that ensuring consistency, completeness, and accuracy of the data requires a rigid collection protocol. For example, in the Academic Tree dataset individuals may use all uppercase letters or put a nickname in parentheses, all of which create problems for automated analysis. A unique numerical identifier would allow for much less variation. This problem is clear to many in the scientific community, and it is being pursued by projects such as ORCID ^25^. However, to facilitate performance analyses, its use must be required, especially in the authorship section of papers, so that the publications for authors with the same name can be easily distinguished in an automated fashion.

## Conclusion

Using methods of network analysis, Nobel laureates were identified as a highly connected group in The Academic Tree network. Several successful mentoring communities could be identified using the number of Nobel laureates as a measure of scientific success. A variety of performance measures exist that would increase the sensitivity of these types of analyses and would allow for the exploration of a greater variety of questions if the measures were collected and made available in a single database. This could provide valuable information regarding individual, institutional, and national factors associated with success in modern science and lead to a greater understanding of best practices. The rewards in such an endeavor would be large, especially in the current climate where there is an increased focus on effective collaboration and teamwork.

## Methods

### Network Collection.

The Academic Tree Network of mentor-mentee relationships was obtained for analysis from Academic Tree on November 2, 2015 ^7,8^. Academic Tree is a web-based database of academic mentor-mentee relationships that uses a crowd-sourcing method for the collection of information. Individuals can voluntarily provide information regarding academic relationships through the Academic Tree website. Academic Tree can be decomposed into an interconnected set of 68 domain specific networks, and it is possible for an individual to be listed in more than one domain. For example, a Cell Biology Tree exists for individuals working in cell biology, and a Genetics Tree exists for those working in the field of genetics. An individual working in both areas can identify themselves as belonging to both trees.

As can be seen in Table S2, the Academic Tree database holds several types of information, including an individual’s specific research area, major research area (i.e., one or more of the domain specific trees), and five possible academic relationships between individuals in the network, including doctoral student-advisor relationships. The database was received from Academic Tree in SQL format which included an edge file and a node file. There were 114,949 entries in the node file and 260,201 entries in the edge file.

### Network Filtering.

Several steps were taken to enhance and filter the network prior to analysis. In the original files, there were 484 individuals listed as Nobel laureates. However, 47 additional Nobel Prize winners could be identified in the file and were labeled as such. The Nobel Prize category and year were added for all Nobel laureates. All information regarding Nobel laureates was obtained from the Nobel Foundation (1). This network was first filtered to include only relationships between doctoral students and advisors (see Table S3, Filter 1). Next, the network was filtered to included only individuals listed in at least one science tree (Table S3, Filter 2). As a result, the majority of Nobel laureates winning prizes for peace, literature, and economics were removed. In the last step, the strongly connected components within the larger network data set were identified using network analysis tools in Gephi ^26^. Table S4 lists the number of strongly connected components of different sizes along with the number of nodes and the number of Nobel laureates associated with each component size. The largest strongly connected component of 57,831 nodes was significantly larger than any of the other components and held the vast majority of Nobel laureates (402 of 472). In fact, in Table S5, this was a significant majority of all Nobel laureates in physics (58.2 %), physiology (65.2 %), and chemistry (86 %), justifying the use of this subnetwork in the subsequent analysis. All Nobel laureates in the largest strongly connected component of the network received prizes in chemistry, physics, and physiology or medicine with one exception, Herbert Simon, a highly interdisciplinary scientist who won the Nobel Prize in economics. There were no prize winners in literature, and only one Peace Prize winner, Linus Pauling, who was also awarded the Nobel Prize in chemistry. All further network visualization and filtering was done in Cytoscape ^27^. The Cytoscape filtering tool was used to identify and visualize the subnetworks displayed in Fig. 1B, Fig. 2, and Fig. 3.

### Data Analysis.

A breadth first search algorithm was used to calculate the number of family members and the number of Nobel laureate family members for each individual. This was instantiated in a custom C++ program which takes a directed acyclic graph as a node and edge list. The node list contains node id, Nobel status, and Nobel Prize category. The edge list contains source and target nodes. The direction in the network (forward, backward, or both) and the number of academic generations (i.e., steps in the network) to be calculated is specified as input to the program. Number of ancestors/Nobel ancestors was calculated as 31 steps (i.e., the diameter of the network) backward while number of descendants/Nobel descendants was calculated as 31 steps forward. The number of mentees/grandmentees and Nobel mentees/grandmentees was calculated as two step forward in the network, and the number of local family/Nobel local family was calculated as three steps forward and backward in the network. While calculating number of Nobel laureate family the program also tracks number of Nobel Prizes in each category.

### Heterogeneity Computation.

A measure of heterogeneity was used to calculate the diversity of Nobel Prizes awarded within three steps of each individual in the network (Equation 1) where *L* is the number of Nobel laureates within a specified distance in the network, *N* is the number of Prize categories (5 in this case), *n_i_* is the number of Nobel laureates in a specific prize category within a specified distance. The number of Nobel laureates in an individual’s local family was an essential factor in the equation and meant that the scores could not be compared across individuals with different numbers of Nobel family members. Therefore, the scores were normalized to fall between 0 and 1 with 1 representing the greatest possible diversity for a given number of Nobel laureates.

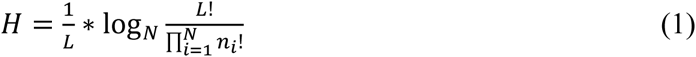

### Generation of Random Networks

Hypothesis testing with network data is problematic in that the assumption of independent observations required for many statistical methods is violated, resulting in standard errors that are computed incorrectly ^28,29^. To handle this, the significance levels in our analyses were adjusted by creating a distribution of expected test statistics, derived from random samples, for comparison with an observed test statistic ^29^. This involved permuting values for the predictor variable (Nobel status) with respect to an outcome variable (number of Nobel laureate family members) for one thousand samples, performing the statistical analysis on each of the random samples, and then counting the number of test statistics on the permuted data that were greater than or equal to the observed statistic. This number was then divided by the number of random samples to produce an adjusted *p* value. For example, if three random test statistics of 1000 permuted samples are greater than or equal to the observed test statistics, the *p* value would be adjusted to 0.003.

To accomplish this, the C++ program described earlier had options available for generating 1,000 networks with Nobel status randomly assigned to nodes across the network in the same proportion as the true data, each time recomputing outcome measures for each node. As can be seen in Fig. 4, this produced alternate networks with equivalent topology (i.e., the same number of family members and academic structure for each node) but randomly distributed Nobel laureates and thus, random outcomes.

**Fig. 4.**
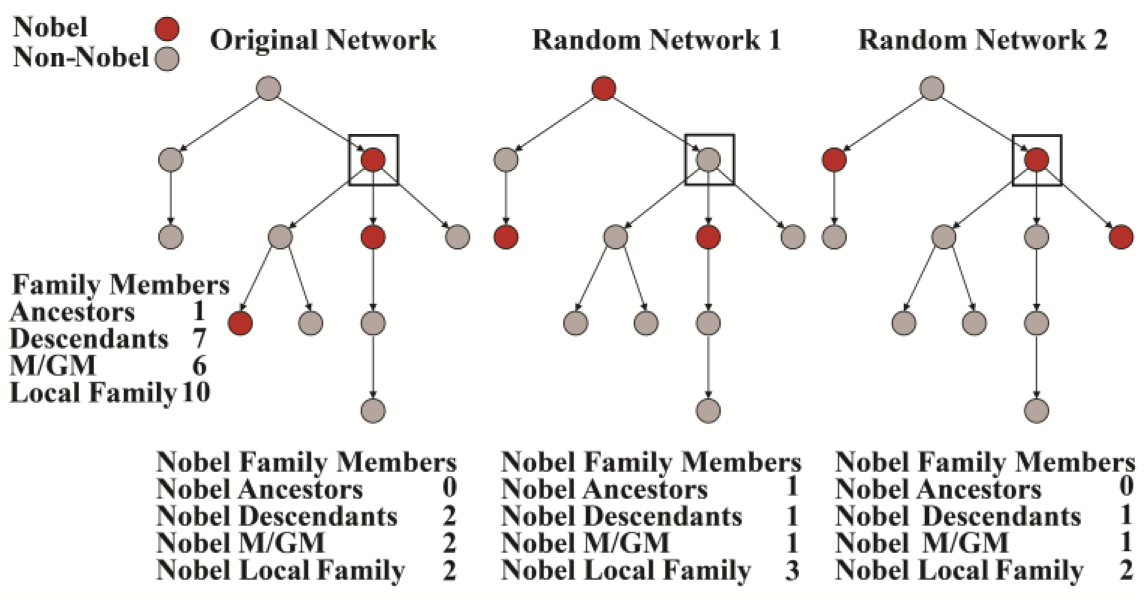
Number of family members and number of Nobel laureate family members computed for an individual network node, highlighted in the box, for a directed network (left) and two networks in which Nobel status is randomly permuted (right).

## Acknowledgments

The authors would like to thank Stephen David for providing The Academic Tree database. This research would not have been possible without his generous support. The authors would also like to thank anonymous reviewers and members of the KBRIN Bioinformatics Core for helpful insight and feedback. All data and scripts used in constructing this analysis is available at http://bioinformatics.louisville.edu/Nobel/. ECR conceived the idea of the project and supervised all aspects of the project. JHC implemented the computational aspects of the project, updated and corrected the mentor network, and performed all network analyses, including biographical and historical reviews. JRP provided insight into the analyses from a social science perspective and helped with preparation of the manuscript. YZ implemented the web interface for interactively exploring the Nobel networks. All authors contributed to the writing of the manuscript. Support for JHC and ECR provided by National Institutes of Health (NIH) grant P20GM103436 (Nigel Cooper, PI). The contents of this work are solely the responsibility of the authors and do not represent the official views of the NIH or the National Institute for General Medical Sciences (NIGMS).

